# Modelling structural determinants of ventilation heterogeneity: a perturbative approach

**DOI:** 10.1101/329961

**Authors:** Carl A. Whitfield, Alex Horsley, Oliver E. Jensen

## Abstract

We have developed a computational model of gas mixing and ventilation in the human lung represented as a bifurcating network. We have simulated multiple-breath washout (MBW), a clinical test for measuring ventilation heterogeneity (VH) in patients with obstructive lung conditions. By applying airway constrictions inter-regionally, we have predicted the response of MBW indices to obstructions and found that they detect a narrow range of severe constrictions that reduce airway radius to 10%–30% of healthy values. These results help to explain the success of the MBW test to distinguish obstructive lung conditions from healthy controls. Further, we have used a perturbative approach to account for intra-regional airway heterogeneity that avoids modelling each airway individually. We have found, for random airway heterogeneity, that the variance in MBW indices is greater when indices are already elevated due to constrictions. By quantifying this effect, we have shown that variability in lung structure and mechanical properties alone can lead to clinically significant variability in LCI and *S*_cond_, but only in cases simulating obstructive lung conditions. This method is a computationally efficient way to probe the lung’s sensitivity to structural changes, and to quantify uncertainty in predictions due to random variations in lung mechanical and structural properties.

## Introduction

The relationship between structure and function in the human lung is an important research area in physiology and medicine. The structural changes associated with various obstructive lung diseases, such as cystic fibrosis (CF) and asthma, can give rise to ventilation heterogeneity (VH) where inhaled gas is unevenly distributed in the lung, leading to poorer gas mixing efficiency [1,2]. The severity of these conditions, in particular CF, is often quantified clinically using Multiple Breath Washout (MBW) tests [3]. MBW uses an inert tracer gas to quantify how effectively fresh air is turned over in the lung by measuring the tracer gas concentration and flow rate at the mouth. These data are used to compute clinically tested indices such as the lung clearance index (LCI) and phase-III slopes [4], which are indicators of VH.

Modelling the results of MBW tests accurately is challenging. First, modelling gas flows in a heterogeneous lung structure is a computationally expensive task. According to cast estimates there are (on the order of) 10^4^ – 10^5^ conducting airways (those where no gas exchange occurs) [5,6], but on the order of 10^7^ branches in total including the acinar ducts (in the alveolar region of the lung). There have been numerous approaches to resolve this problem, such as using a symmetric airway network with an effective diffusion coefficient to account for heterogeneous ventilation [7], compartmental models with asynchronous or asymmetric ventilation [8-10], modelling a single heterogeneous acinus [11,12], and replacing the acini with well mixed units or symmetric models [13-17]. Second, many model parameters are difficult to measure experimentally, and are variable between subjects, which increases the uncertainty in model predictions. Quantifying this uncertainty is a key step to making more clinically relevant model predictions.

In this paper, we first introduce a reduced model of lung structure that accounts for asymmetry in the inter-lobar airways and inter-regional heterogeneity but assumes symmetrical branching in the lower airways. We label this ‘model M’. In the baseline case, all airways in a given generation of each lung region are assumed identical, so that each region can be represented by a single ‘mean-path’. To simulate the effects of obstructive lung disease we systematically apply constrictions to the lower airways in model M, which gives rise to inter-regional VH. This model is only an approximation to the true lung mechanics, assuming linear elastic response of the alveoli and Poiseuille flow throughout.

Second, we outline a perturbative approach that extends ‘model M’ to include weak intra-regional heterogeneity. This method estimates the effect of changes in mechanical or geometrical properties of the airway tree within the symmetrically-branching regions. The linearised responses of the model outcomes to these perturbations are then superposed to reconstruct the full airway tree, at low computational cost. We label this perturbative method ‘model P’. Model P can be used to describe lung structures with heterogeneity that is either deterministic (where the structural and mechanical properties are prescribed) or probabilistic (with properties described by multivariate distributions) using the same simulations. The probabilistic descriptions of lung structure means that we are able to directly estimate the variance of model outputs due to the parameter distributions and thus quantify uncertainty. We have applied this method to the particular case of MBW simulation in “Model P: Intra-regional heterogeneity”.

The aim of this research is to quantify the sensitivity of MBW indices to structural heterogeneity for both healthy and diseased lung models. The perturbative method developed makes it tractable to probe the geometry of all airways (including acinar ducts) efficiently and relate them directly to MBW outcomes and the distribution of ventilation in the lung.

## Methods

### Model M: Ventilation and transport in symmetrically branching lobar regions

We have modelled the effects of inter-regional heterogeneity in the lung using a coupled network model of ventilation and gas transport in the airways and acinar ducts. To reduce model complexity, we have initially assumed that the airway tree can be approximated as completely symmetric within seven lung regions (see Figure 1); we refer to this as model M. Each region corresponds to a lobe or lobar compartment resembling the regions in Horsfield and Cumming’s lung model M [6]. We define the Strahler order of an airway by the maximum number of generations distal to it including acinar duct generations (counting from zero). Details of the model and parameters used can be found in S1 File and are summarised in S1 Table.

**Figure 1:**
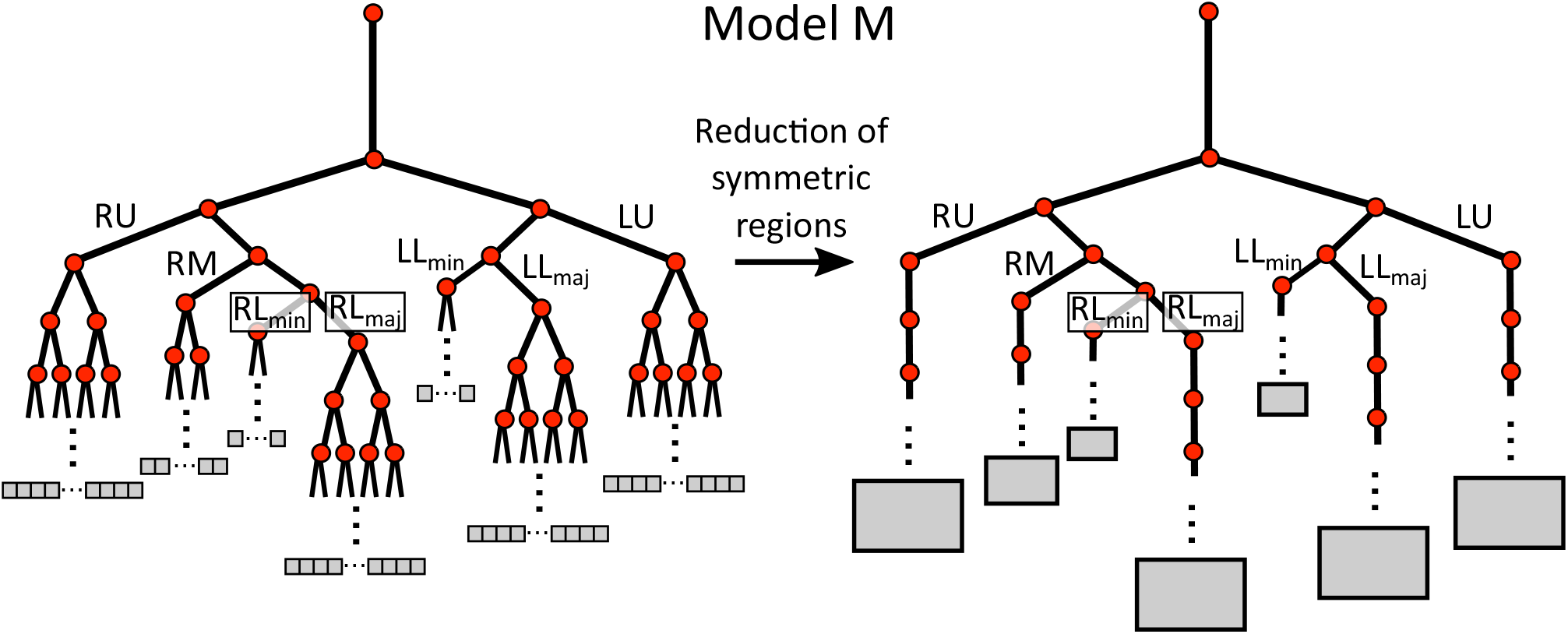
Network diagram of model M used to simulate MBW. Each lobar region (right-upper, RU; right-middle, RM; right-lower minor, RL_min_, right-lower major, RL_maj_; left-upper, LU; left-lower minor, LL_min_; left-lower major, LL_maj_) is modelled a symmetrically branching tree (left) which can be modelled as a single path (right). Each black line represents an edge of the network, while each red dot is a vertex. The grey boxes indicate the parenchymal volume fed by each terminal airway or set of airways.

Gas flow on the airway network is calculated using a coupled set of ordinary differential equations (ODEs) that account for airway resistance (assuming Poiseuille flow), linear viscoelasticity of the acinar units and a uniform applied pleural pressure (similar to [14,15,19]),

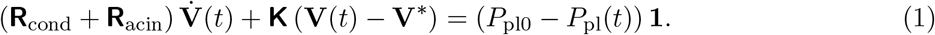

The vector **V** contains all the lung unit volumes (represented as grey boxes in Figure 1). The tensors **R**_acin_ and **K** are diagonal and contain the resistance and elastance of the lung units respectively. The full tensor **R**_cond_ is the airway resistance matrix, defined in S1 File §1.2 and the scalar *P*_pl_(*t*) is the constant pleural pressure. The vector **V**\* corresponds to the resting volumes of the lung units at zero flow where *P*_pl_(*t*) = *P*_p10_. These ODEs are solved directly using the Eigen [20] factorisation routine ‘PartialPivLU’ in C++, as outlined in detail in S1 File §1.2.

The concentration of inert gas on the network is then calculated using a one-dimensional advection-diffusion equation (S1 File §1.3) that accounts for transport into the alveolar sacs (similar to [11,12,21]). On a given edge *e_i_*, which can represent a single airway or numerous identical airways in the same generation, the transport equation is

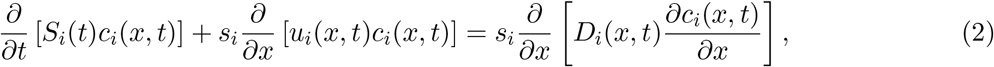

where *s_i_* and *S_i_* are the inner and outer total airway cross-sections in *e_i_*. For conducting airways *s_i_* = *S_i_*, and in the acinar ducts within the lung units *S_i_* – *s_i_* accounts for the (time-dependent) volume of the acinar sacs lining the ducts. This is essentially the ‘trumpet’ representation of [22] and is outlined in more detail in S1 File §1.4. The effective diffusion constant *D_i_*(*x,t*) is given by Taylor-like dispersion described in [23] in the conducting airways,

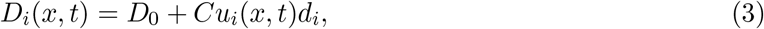

where *d_i_* is the airway diameter and the constant *C* is defined as *C* = 1.08 for inspiration and *C* = 0.37 for expiration. In the acinar ducts, pure molecular diffusion is modelled but with a modified airway cross-section, as described in [12],

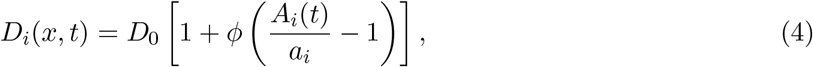

where *ϕ* is a phenomenological parameter that sets the fraction of alveolar sac cross-section that is involved in diffusion. However, this relation by no means captures the full complexities of gas mixing dynamics in the pulmonary acinus (*e.g*. [24-26]). The resulting transport partial differential equations (PDEs) are discretised using a finite volume method (detailed in S1 File §2), and solved iteratively using the Eigen [20] ‘BiCGSTAB’ routine.

To simulate MBW in model M the concentration is initialised to *c* = 1 everywhere in the lung. Then a regular sinusoidal breathing cycle is simulated with *c* = 0 at the mouth on inhalation, and a diffusive-flux-free *∂_x_c* = 0 boundary condition on exhalation. Parameter values for tidal volume (*V_T_*), functional residual capacity (FRC), airway dead-space volume (*V_D_*) and lung elasticity (*K*_lung_) are chosen to be representative of a healthy adult male (see S1 Table).

The MBW test measures FRC from the total inert gas exhaled over the test [4], which we label as 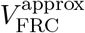. The lung clearance index (LCI) value is the number of lung turnovers (exhaled volume in units of 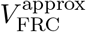) required to reduce the concentration (measured at the mouth at end of exhalation) to 2.5% of its initial value. In this paper we interpolate this number of lung turnovers to measure small changes in LCI that are below the resolution of clinical LCI measurements (see S1 File §3). LCI is a widely tested clinical measure of VH [27,28], with healthy values generally in the range 6–8 and larger values indicating increased heterogeneity. Phase-III indices measure VH through the slope of inert gas concentration versus volume for individual exhalations, and their interpretation is informed by numerical modelling [29–31]. In this paper we focus on *S*_cond_, the linear gradient of the normalised phase III slopes, measured according to the clinical guidelines for MBW in [4].

To measure VH in the model directly, we calculate the fractional ventilation (FV) of the lung acini, which is the dilution rate of the inert gas concentration between successive breaths at end-tidal volume. Hyperpolarised helium MRI imaging [32] is a state-of-the-art technique that can be used to measure the FV distribution in the lung [33]. Heterogeneity in FV correlates strongly with increased LCI, but the MRI images can also identify changes not picked up by MBW indices, as well as provide important information about the spatial distribution of FV [3]. The network model presented here cannot recover the spatial distribution of FV measured in MRI, but does predict the probability distribution of FV in the lung.

### Model P: Perturbative method and application to structural heterogeneity

Building on model M, we use perturbation theory to calculate the changes in gas concentration due to small variations of the properties of a single airway or acinus in a symmetrically-branching lung region. Simulation variables, such as LCI, are complicated non-linear functions of the input parameters. Here, we restrict our attention to variations in airway cross-section and length, and acinus elastance, parametrised by the dimensionless 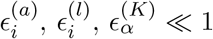 respectively, where *i* is the airway index and *α* the acinus index. LCI, for example, is then reconstructed by superposition of the linear responses,

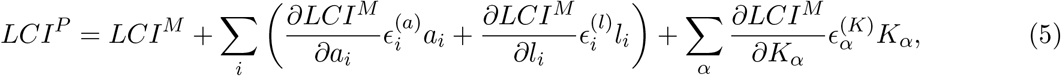

where the *M* and *P* superscripts refer to variables in model M and P respectively, and the sum is taken over all perturbed airways or acini in the model. We define the linear sensitivity of variable *g* to parameter *f* as

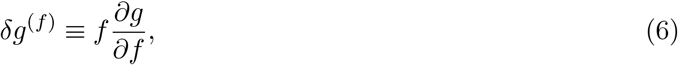

enabling the comparison of sensitivities with respect to different parameters. Many of the linear sensitivities in (5) are degenerate due to the symmetric nature of the regions in model M, and so reconstructing the whole solution requires simulating only the linear response to one perturbation for each generation of each region (see Figure 2), and thus the system size grows linearly with the number of generations included, rather than exponentially. This is outlined in more detail in S1 File §1.5.

In “Model P: Intra-regional heterogeneity”, we use the linear sensitivities computed for model P to calculate the variance of MBW indices due to intra-regional heterogeneity. These sensitivities are computed for realisations of model M covering a range of constriction severity. In the simple case where the perturbations are modelled as variables in a multivariate Gaussian, the variance in model outputs can be approximated as a sum of the covariance of inputs weighted by their respective linear sensitivities

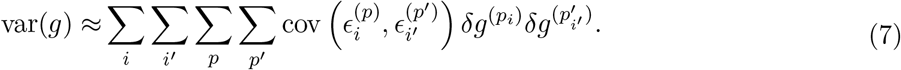

In eq. (7), the sums are taken over all perturbed airways labelled by the indices *i* and *i*’, where *p* and *p*’ refer to the corresponding perturbation (*a* or *l*; we disregard perturbations to *K* as we find that airway geometry dominates the variance on the MBW outcomes). The variable *g* can refer to any property in the model, and explicit examples are given in S1 File §3. This gives a computationally efficient method to relate weak intra-regional heterogeneity to variance of model outputs that relies only on each constricted case being simulated once.

To construct the multi-variate Gaussian airway distributions, we assume that the coefficients of variation of the airway areas and lengths, *σ_a_* and *σ_l_* respectively, is independent of generation. First, we model perturbations to be independently normally distributed

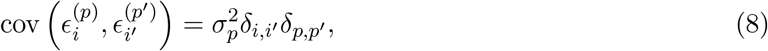

where *δ_xy_* is the Dirac delta function. Second, we assume that the perturbations are normally distributed around the perturbation to their parent branch, and that area and length perturbations in the same branch are correlated with coefficient *ρ_al_* such that

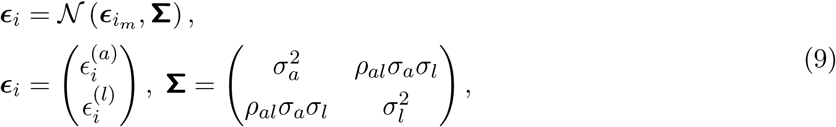

where *i_m_* is the index of the parent branch to airway *i*. This case results in what we term ‘structurally correlated’ heterogeneity within each symmetrically-branching lung region. This means that airways and acini that are closely related on the tree have more closely correlated fluctuations in airway geometry (see Figure 3). This is similar to the random structure generated used for particle deposition calculations in [34], whereby airway sizes are generated according to the size of the parent airway.

**Figure 2:**
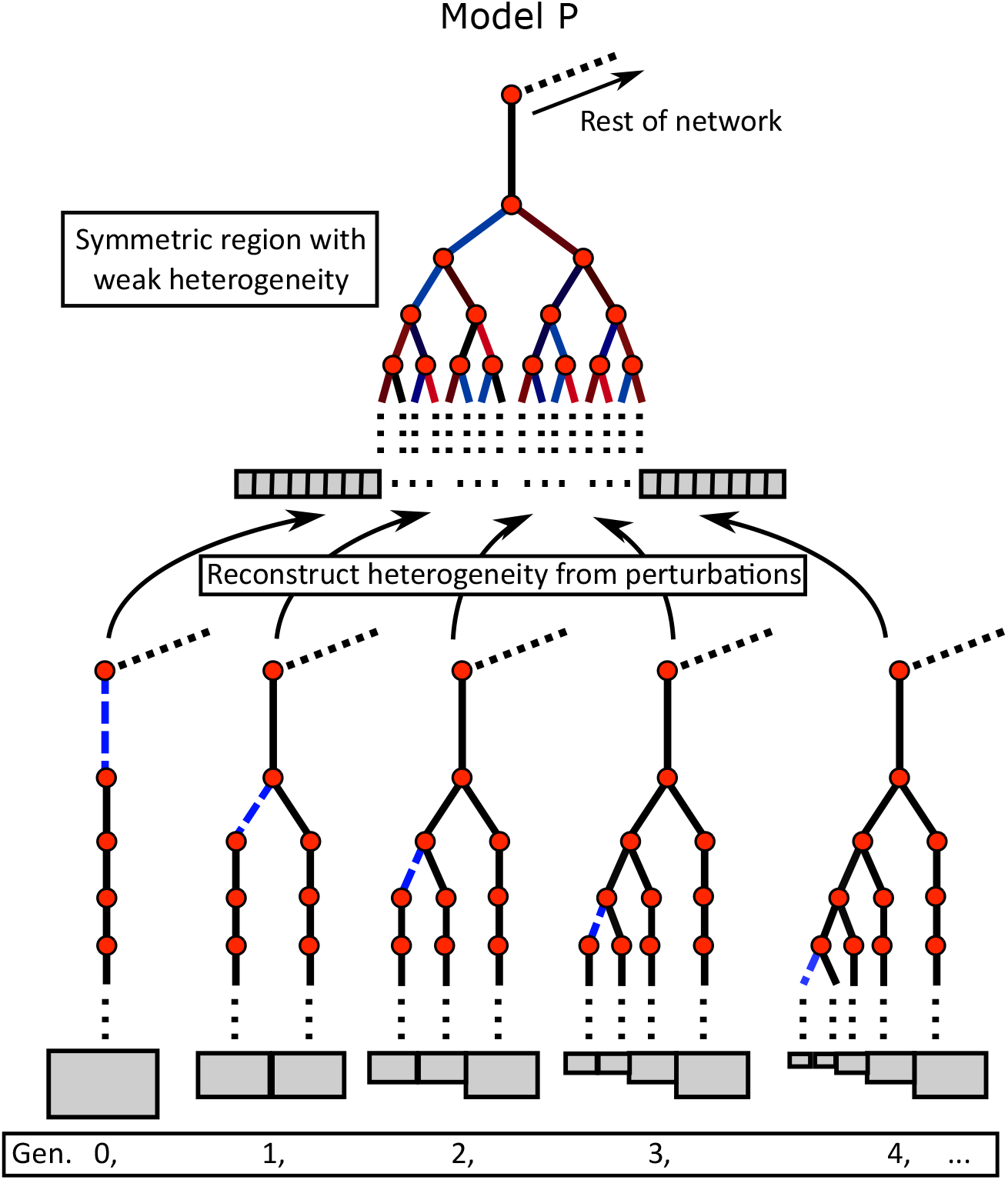
Top: Sketch of the airway network within a symmetric lobar region, the heterogeneous colouring of the edges represent small changes in geometry away from complete symmetry of model M. This weak heterogeneity can be reconstructed using the sensitivities computed from a single perturbation to each perturbed property in each generation added to the initial model M solution. Bottom: The airway networks used to compute the linear sensitivities corresponding to each generation as labelled, where the blue dashed edge is the perturbed airway.

**Figure 3:**
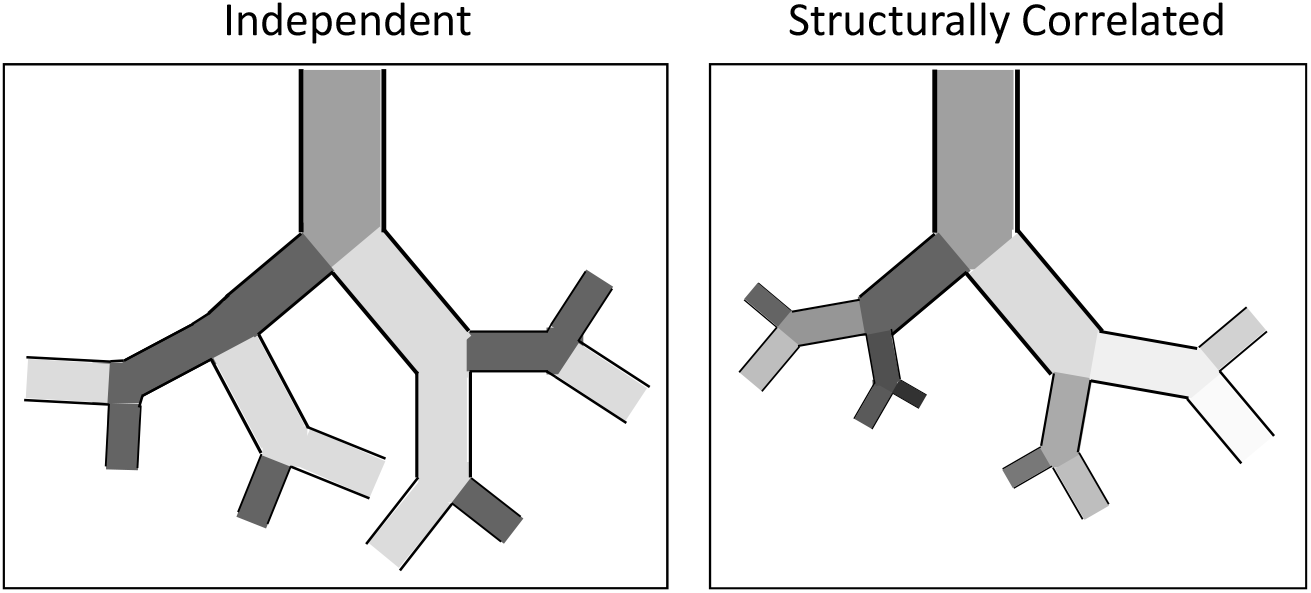
Sketch of randomly distributed airway sizes, shading (from dark to light) indicates airway size relative to generational average. In the independent case, variations in branch size are uncorrelated. In the correlated case larger than average airways are likely to beget airways that are also larger than average, resulting in an increased uncertainty in the size of the most distal airways. For simplicity, all airways in this sketch have fixed aspect ratio (the case *ρ_al_* = 1). When *ρ_al_* = 0, airway length and area are independent.

Finally, we use these results to compute the probability density of any acinus in the model having a given FV value (averaged over the whole MBW test). In the limit of a large number of acini, the distribution of FV values in any given lung model realisation will tend towards this distribution. For further details of the airway heterogeneity models used and the method of reconstructing the FV distributions see S1 File §3.

### Model validation

Validation studies to test the accuracy and precision of the numerical simulations can be found accompanying the source code at [35]. In these we have tested that the code is suitably converged for the choice of model time-step and space-step, and that inert gas volume is conserved. We also tested that the perturbative model converges exactly to the mean-path model in the limit of small perturbations.

We have validated the predictions of model M by comparing the outcome to predictions of interregional heterogeneity in a simple analytical two-component model in S1 File §5. To validate model P, we ran Monte Carlo simulations on a version of model M with some intra-regional heterogeneity and compared the measured variance in LCI to that predicted by model P (these results are presented in S1 Figure).

## Results

### Model M

The healthy baseline case of model M (no airways constricted) assumes that all regions have the same airway sizes and mechanical properties. This results in a homogeneous distribution of gas, with the only asymmetry originating from the arrangement of the proximal airways supplying the lobar regions, and the number of generations within each region. In this case the simulations show little regional difference in FV and each lobe contributes proportionately to the washout. The baseline values of the MBW indices (using SF_6_ as the tracer gas) were LCI = 5.16, *S*_cond_ = 1.92 × 10^-4^L^-1^ and 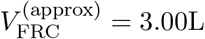 (to 3 s.f.). Using N_2_, the LCI reduced slightly to 5.04, due to better gas mixing and a more proximal diffusion front (note that the effects of gas exchange are not included in this model).

### Model M: Inter-regional heterogeneity

Figure 4 (solid lines) shows the effect of constricting airways in the right-middle (RM) lobe of model M at three different depths. Each case simulates localised bronchoconstriction, with all airways in a given generation range (proximal, central or distal) reduced in cross-section by the same fraction, approximating the pathophysiology of asthma or CF. A marked response in all three MBW indices is evident for radius constrictions above circa 70%. The responses of LCI and *S*_cond_ to airway constrictions are strongly correlated, peaking at approximately 80% constriction of the radius before dropping back to baseline values (Figures 4(a) and (b)).

**Figure 4:**
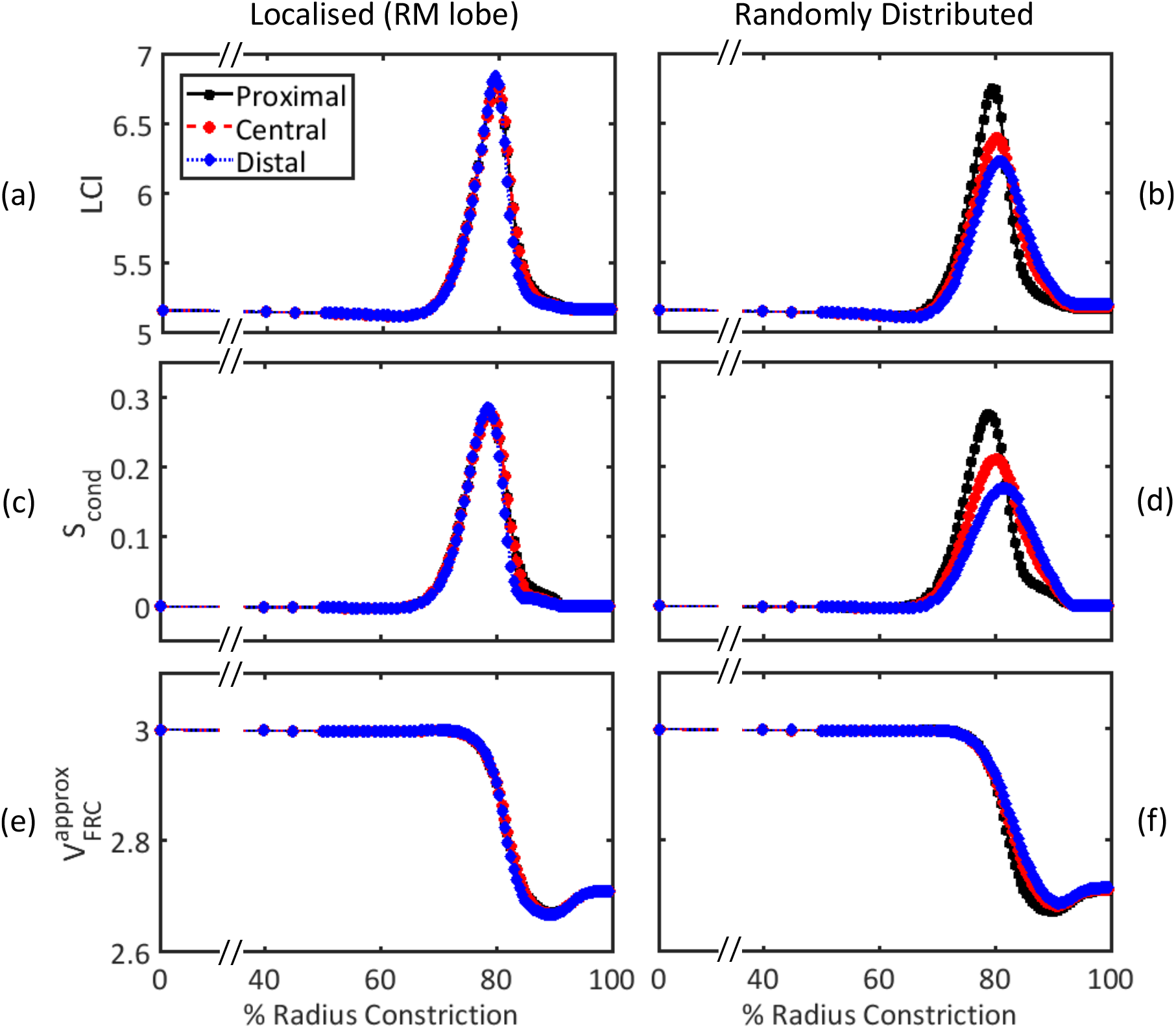
Relationship between constriction strength (% radius reduction) and MBW indices simulated using model M. (a)-(b) LCI, (c)-(d) *S*_cond_ and (e)-(f) measured FRC volume 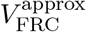 with constrictions applied to airways feeding 10% of the lung acinar volume. Three different depths were tested, corresponding to Strahler order 19-16 (proximal, black squares), 15-12 (central, red circles), and 11-8 (distal, blue diamonds), where all branches constricted were taken to be directly descended from the most proximal in all cases. (a), (c) and (e) show simulations where constrictions were all localised within the right-middle lobe, whereas (b), (d), and (f) show realisations where the positions of the constrictions were uniformly randomly distributed throughout the lung. Example animations of localised and random constrictions are shown in S1 Video and S2 Video respectively.

Simulated LCI and *S*_cond_ are effectively independent of constriction depth and drop off at larger constrictions, where the constricted region becomes essentially unventilated and thus undetectable at the mouth. This is shown in Figure 4(e) by the reduction in measured FRC volume of approximately 10% (295 ml) of the lung volume. This compares well to the simplified analytical prediction of VH in a two-component model (S1 File §5).

It is a simplification to assume that constrictions or blockages would be localised to a single lobe. However, randomly distributed constrictions applied to families of airways at each depth that feed the same fraction (10%) of the lung volume, result in a very similar response due to the homogeneity of the baseline case (see Figures 4(b), (d) and (f)). The response is weaker than the localised case for a more distal heterogeneous distribution of constrictions, and drops off more gradually at >80% radius constriction.

To summarise, we have found that MBW indices detect a restricted range of severe airway constrictions, which our results predict to be most sensitive when airways are between 10% - 30% of their original radius.

### Model P

Model P consists of simulating the linear difference in all model variables due to individual perturbations to airway geometry or acinar elastance. These linear sensitivities are then combined to predict how sensitive the outcomes of model M are to weak intra-regional heterogeneity and changes in global model parameters.

### Model P: Intra-regional heterogeneity

Figure 5 shows the predicted standard deviation in MBW indices due to a random distribution of airway geometries and acinar elastances in model P. Two types of random heterogeneity are presented: first where changes to airway geometry are independently normally distributed within the lung regions; and second where they are structurally correlated (see Figure 3). In general, the standard deviations of LCI and *S*_cond_ increased when perturbations were correlated with their parent branch. Note that the mean value of LCI and *S*_cond_, for varying constriction magnitude in the RM lobe, are unchanged from the predictions of model M, since model P only incorporates linear responses. Nonetheless, the approximation of variance remains good for *σ_a_*, *σ_l_* < 0.25 (see S1 Figure).

**Figure 5:**
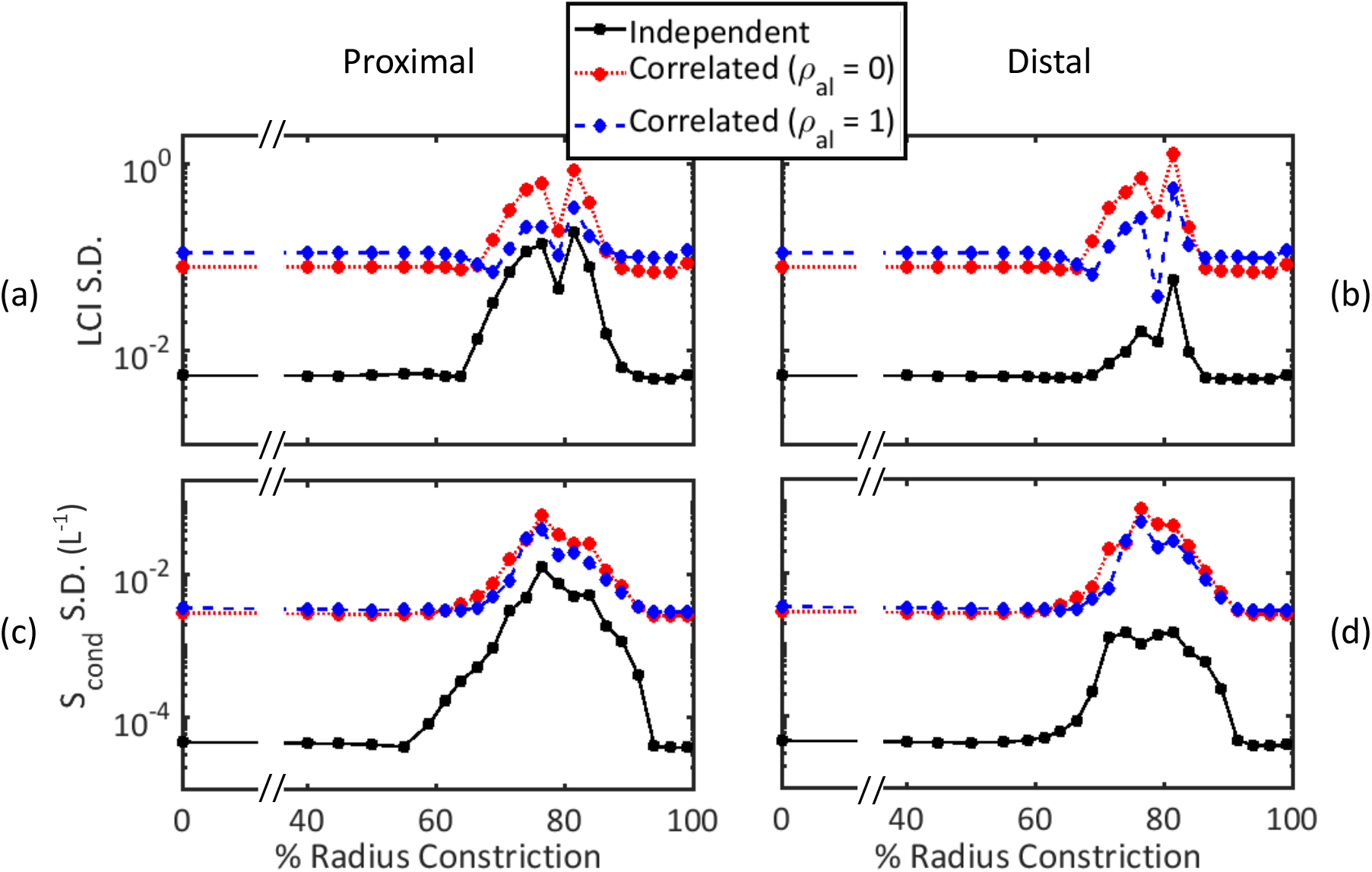
Standard deviation in (a)-(b) LCI and (c)-(d) *S*_cond_ vs. constriction strength (for constrictions confined to RM lobe) predicted using model P. Results are shown for independent normally distributed perturbations (black squares) and structurally correlated perturbations (see S1 File §4.2) with *ρ_al_* = 0 (diameter and length uncorrelated, red circles) and *ρ_al_* = 1 (fixed diameter-length ratio, blue diamonds). (a) and (c) show the results for Strahler orders 19-16 (proximal) and (b) and (d) for Strahler orders 11-8 (distal, dotted line, diamonds). Constrictions were applied to all airways in the RM lobe within these generation ranges as in Figure 4. In all cases *ρ_a_* = 0.2 and *σ_l_* = 0.1. Note the logarithmic scale on the vertical axes.

The standard deviations of LCI and *S*_cond_ due to airway heterogeneity are orders of magnitude larger when the indices are already elevated by constrictions in the RM lobe. There is a small drop in LCI variance at ~ 80% radius constrictions, corresponding to the stationary point of the LCI curve in Figure 4(a)-(b). This means that the uncertainty shows similar behaviour to the magnitude of the gradient of the curve in Figure 4(a)-(b).

Figure 5 also shows different responses depending on constriction depth. Independently distributed airway heterogeneity has less effect on the standard deviations of the indices when constrictions are more distal. However, when parent-daughter airway sizes are correlated, the effect on LCI and *S*_cond_ standard deviations is similar regardless of constriction depth. This implies that this type of structural heterogeneity, whereby uncertainty in airway size grows with each generation, enhances the contribution of the smaller airways to the overall uncertainty.

Finally, Figure 5 includes the structurally correlated case where airways retain a fixed length-to-diameter ratio to linear order (*ρ_al_* = 1). In this case, LCI and *S*_cond_ variances do not increase as much at severe constriction strengths as the *ρ_al_* = 0 case.

To conclude, when the MBW indices are elevated due to severe airway constrictions, they are more sensitive to heterogeneity in airway size. This results in an increase in the variances of MBW indices when airway geometries are randomly distributed.

### Model P: Fractional ventilation distributions

Figure 6 shows the distribution of acinar FV values at different constriction strengths, and for different distributions of airway heterogeneity. As FV in the RM lobe decreases (and LCI increases) the distribution of FV in this lobe also broadens (Figures 6(b) and (c)), before narrowing again at very large constrictions (Figure 6(d)). This shows that the local FV is most sensitive to airway heterogeneity in the same constriction strength range as the MBW indices. When area and length perturbations are correlated (*ρ_al_* = 1) the FV distribution is narrower in the constricted lobe (RM), where airway resistance dominates, and broader in the other regions, where airway dead-space volume is the dominant factor. This clearly demonstrates a link between the width of the FV distribution and the resulting model uncertainty in MBW indices due to airway heterogeneity.

**Figure 6:**
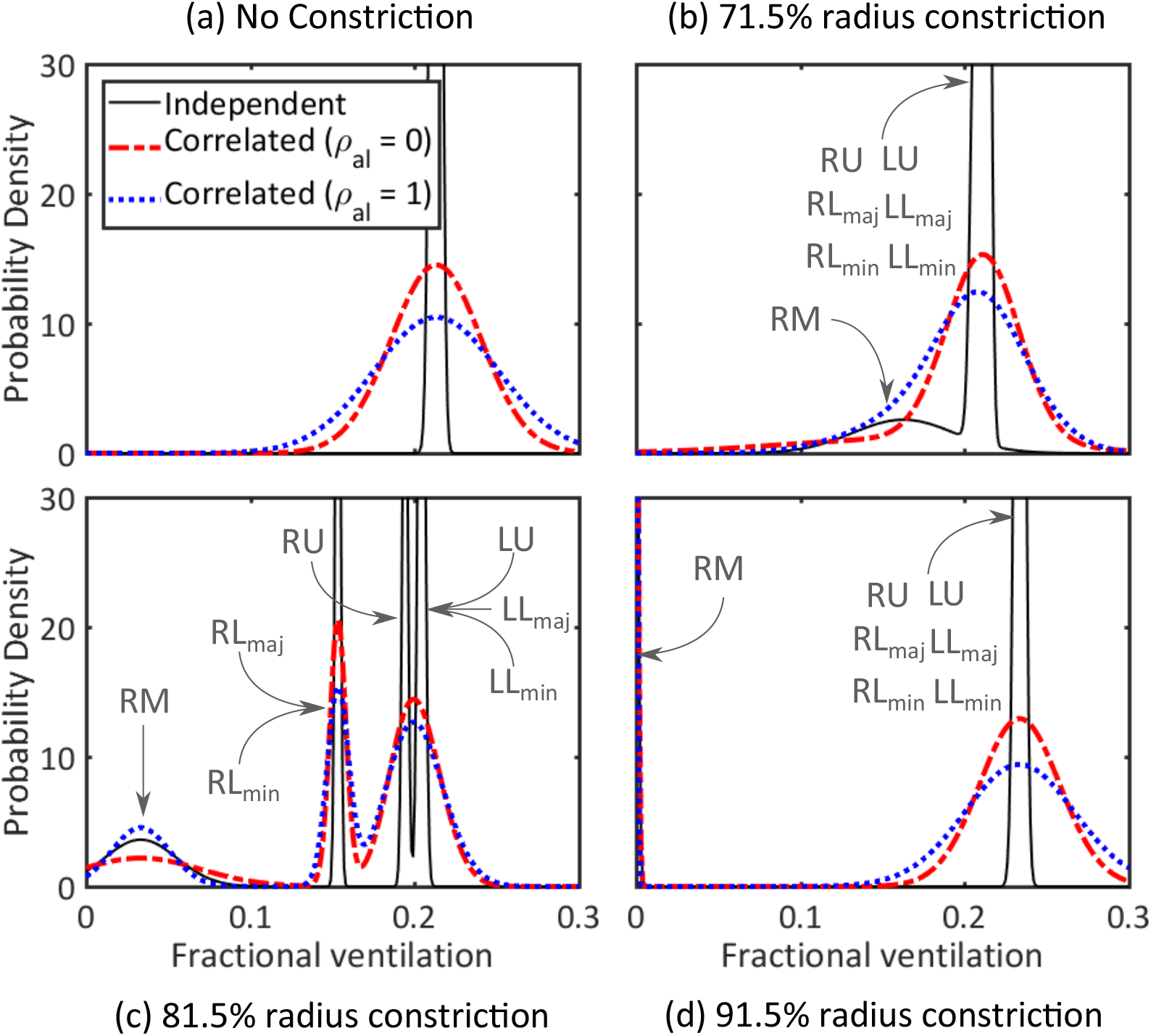
Whole lung FV distributions for (a) 0% (b) 71.5% (c) 81.5%, and (d) 91.5% constrictions to the radius of the airways in the central airways of the RM lobe (central, Strahler orders 15-12) from model P. Results for independent random perturbations (solid black lines) and structurally correlated random perturbations with *ρ_al_* = 0 (dashed red lines) and *ρ_al_* = 1 (fixed diameter-length ratio, dotted blue lines). *σ_a_* = 0.2 and *σ_l_* = 0.1 was used in all cases. In (b), (c) and (d) the peaks are labelled by their corresponding region(s) as denoted in Figure 1. For visibility, the y-axis range does not extend to include all of the peaks.

A key finding is that the unconstricted mean-paths are relatively unaffected, and remain fairly insensitive to airway heterogeneity within those paths. Nonetheless, there is a small drop in FV in the right-lower lobe, which can be explained by a pendelluft effect where gas from the right-middle lobe is re-inspired into the right-lower lobe due to the asynchronous nature of the ventilation (see S1 Video).

In summary, the FV distributions show that severe airway constrictions result in a much broader distribution of gas turnover in the affected lung region (assuming some randomness in airway geometry). Aside from the noted pendelluft affects, the distribution of FV in the unaffected lobes remained largely unchanged, highlighting the parallel nature of the airway network structure.

### Model P: Further insights from linear sensitivity analysis

The linear sensitivities computed for model P give an insight into how MBW indices depend on airway properties at different depths. Examples of LCI and *S*_cond_ sensitivity vs. airway generation are shown and discussed in the supplementary figures. S2 Figure compares the sensitivities using N_2_ and SF_6_ tracer gas in the model in the absence of constrictions. It demonstrates that, for the advection-dominated conducting airways the two responses are identical, whereas in the diffusion-dominated acinar ducts, the responses are markedly different due to the difference in molecular diffusivity. S3 Figure shows the linear sensitivity of LCI to airway perturbations when the airways of the RM lobe are already constricted. In this case, the LCI response is completely dominated by the geometry of those airways that are already narrowed and so have high resistance, and the response is markedly larger than the baseline case in S2 Figure(a,b).

In Figure 7 these local sensitivities are combined to predict the sensitivity of LCI to changes in three global model parameters. In the absence of constrictions, LCI is weakly sensitive to airway dead-space and completely insensitive to airway length-diameter ratio and acinar elastance. When constriction magnitude is elevated (to 70–90% of the radius) LCI becomes most strongly sensitivity to length-diameter ratio, and also much more dependent on elastance, while the dependence on total dead-space remains weak. Generally, a very similar response is predicted for *S*_cond_, except that it is not sensitive to *V_D_* for mild constrictions (not shown).

**Figure 7:**
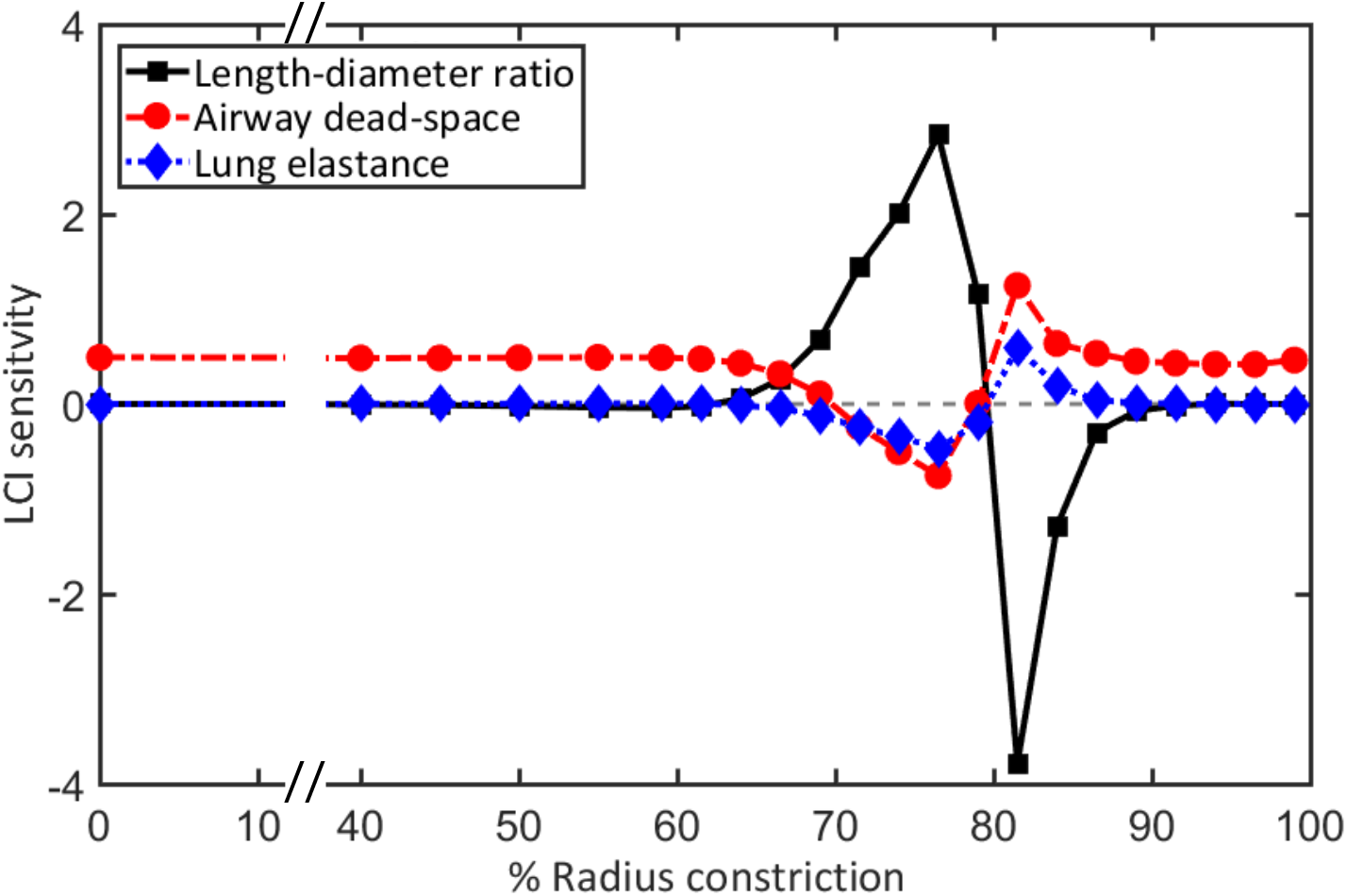
Linear sensitivity (computed for model P) of LCI and to changes in three global model parameters vs. baseline constriction magnitude (of central airways in RM lobe). In the notation of eq. (6) the plot shows *δLCI*^*LD*_cond_^ (length-diameter ratio, black solid line), *δLCI*^*V_D_*^ (airway dead-space, red dash-dotted line), and *δLCI*^*K*_lung_^ (lung elastance, blue dotted line). Each point corresponds to a single simulation of model M with localised constrictions (where the baseline LCI values are given by Figure 4(a)).

Thus, by looking in more detail at the sensitivities computed in model P, we can gain insights into how the MBW indices depend on different sets of airways and global model parameters.

### Model P: Computational efficiency

It is difficult to compute the reduction in simulation time that this method represents as the equivalent non-perturbative calculation involves simulating transport repetitively on all ~ 10^7^ branches, which remains impractical. Nonetheless this is achieved by model P in approximately 5.5 hours on a single processor. If we reduce the calculation to only resolving heterogeneity in the conducting airways (40, 960 terminal branches) then the equivalent non-perturbative calculation takes approximately 12 hours for a single realisation whereas model P takes 0.6 hours under the same conditions (on average). This represents a significant improvement when one considers the number of realisations of the non-perturbative approach required to effectively estimate outcome variance via Monte-Carlo sampling or similar (which only requires a single realisation of model P).

## Discussion

In this paper we have introduced a simple lung model (model M) to account for VH due to structural changes in the airway tree. Our results show that *S*_cond_ and LCI respond in a highly correlated manner to bronchoconstriction, whether this is localised to a single lobe or distributed across the lung (Figure 4). The response is notably non-linear, showing a high sensitivity to large-magnitude constrictions (~ 80% reduction in radius) before returning to baseline values at even larger constrictions. This suggests a mechanism to explain the success of MBW indices to differentiating obstructive lung conditions such as asthma and CF, where airway inflammation and blockage is a typical feature, from healthy volunteers where such narrow airways are unlikely to be present in large numbers.

The sharp response in simulated MBW indices is due to the inverse-fourth-power dependence of airway resistance (approximated here by Poiseuille flow) to changes in airway radius and has also been observed in studies using image-based models with non-linear pressure-drop relations [13,17]. Using a simple two-component model, we show in S1 File §5 that the ratio *r*_DA_/*K*_lung_*τ* (where *r*_DA_ is the baseline resistance of the lower airways) is crucial to determining the constriction strength required to observe an increase in LCI. Literature values predict that the lung elastance greatly outweighs the resistance contribution in healthy lungs, and so we see an increase here only for very severe constrictions.

Model P expands on model M by using perturbation theory to approximate the response of the MBW outcomes to intra-regional heterogeneity. Using this, we found that the uncertainty in predictions due to weak heterogeneity in the structure is greatly amplified when LCI and *S*_cond_ are elevated due to constrictions (for constrictions confined to a single lobe, see Figure 5). This response is dominated by the increased sensitivity to the geometry of those constricted airways in particular. When airway heterogeneity is independently distributed, the variance is greater when there are few proximal constrictions rather than numerous distal constrictions, due to the random contributions averaging out. This uncertainty is amplified further by including structural correlations that account for the inherited nature of airway sizes. We found that the distal airways contribute equally to the variance of MBW indices in this case, because more proximal fluctuations from the mean were propagated down the airway tree.

More broadly, these results suggest that elevated MBW indices induced by airway constrictions are more variable in general, which is observed experimentally in CF and asthma patients through the increased sensitivity of LCI to posture [36,37], as well increased inter-patient variability [38]. Such variability can also be affected by disease severity, as well as randomness in the mechanical and structural properties of the lung considered here.

Furthermore, we have used model P to compute probability distributions of acinar FV values (Figure 6). The predicted FV distributions are generally narrower than measured experimentally [33]. This is in part due to the simplified lung structure and the assumption of weak heterogeneity. Furthermore, acini sizes are also variable which directly affects their FV values, while gravitational effects also play a role [14]. However, the response we observe is indicative of the effects of heterogeneity in airway structure alone, and demonstrate the relationship between variation in structure and the distribution of FV within a lung. We saw that, generally, the FV distribution is much broader in the constricted lung region, which is consistent with the increased variance in MBW indices predicted (Figure 5). These calculations also showed that the unconstricted regions are relatively unaffected by the presence of the blockage, due to the parallel nature of the lung structure.

The LCI values predicted by model M are low compared to those measured experimentally [38], but similar to those simulated in more detailed airway tree models [17,18]. The phase-III slopes are practically zero in the absence of constrictions, whereas in a healthy lung, airway asymmetry and acinar duct asymmetry both contribute to positive slopes [18]. These differences can in part be attributed to the idealised nature of this model, which assumes complete symmetry in the acinar structure. This means that mixing efficiency in the model alveolar zone is better than is likely in reality, and LCI is therefore lower.

Model P addresses the effect of intra-regional airway heterogeneity, which is not present in model M, and is valid only for small deviations of properties from the mean. As a result it misses nonlinear behaviour, which can become dominant at increasing perturbation magnitude. Additionally, the number of trees one has to simulate to compute all of the linear sensitivities increases with the number of symmetrically-branching regions in model M (see Figure 2). Thus there is a balance to be struck between the resolution of model M (i.e. how many symmetrically-branching lung regions are used) being sufficient to simulate realistic VH and computational efficiency.

Other assumptions made in model M are likely to affect predicted MBW outcomes. Most significantly, we have neglected the effects of gravity and posture on inter-regional variation, as well as mechanical coupling of the lung units, which are both predicted to affect FV and MBW indices in simulations [14,39-41] and experiments [37,42,43]. The lack of mechanical coupling means that the predicted asynchrony between lung regions may be exaggerated compared to reality, which could indicate why the range of predicted *S*_cond_ values is notably wider than the increase measured between healthy volunteers and CF patients [45]. Furthermore, air flow has been modelled by the Poiseuille relation in all airways, meaning that the effects of inertia [44] and turbulent flow are neglected. Thus the airway resistance is underestimated, especially in the larger airways, meaning that LCI may become elevated at lower constriction magnitude; however we would not expect this to be a strong effect because as flow rate through the constricted airways falls, their resistance reverts back to Poiseuille. Finally, we have not included the effects of gas exchange on inert gas transport, as it is thought to be negligible (except in the case where nitrogen is used as the MBW tracer gas [46]). The limitations imposed by these assumptions are discussed in more detail in S1 File.

## Conclusion

We have developed a simple, computationally efficient model of gas ventilation and transport in the lung (model M). This has been used to model the relationship between airway constrictions, interregional VH, and MBW indices.

We extended model M by using perturbation theory to measure model sensitivity to airway geometries and acinar elastance. These give an quantitative insight into how the MBW indices depend on the airway properties at different depths and in different lung regions. The linear sensitivities to perturbations form the basis of model P, which accounts for the effect of weak intra-regional heterogeneity. This method has the benefit of being computationally efficient (rather than simulating all airways in the model explicitly) and capable of estimating the variance in model variables using a single simulation (rather than requiring numerous samples).

In future, this approach will be further developed to quantify uncertainty in more realistic lung models that are directly informed by imaging data. The principles outlined here will enable a systematic approach that quantifies uncertainty due to both the intrinsic complexity of lung structure and the additional effects of obstructive lung disease or gravity.

## Acknowledgements

The authors would like to thank Dr Tobias Galla for his contributions to the early stages of this project. Furthermore, we acknowledge fruitful discussions with Prof Jim Wild and Dr Guilhem Collier.

CW was funded by the UK Medical Research Council (MRC) grant number MR/R024944/1. CW and OE acknowledge the UK Engineering and Physical Sciences Research Council (EPSRC) for funding through grant reference EP/K037145/1. AH was funded by a UK National Institute for Health Research (NIHR) Clinician Scientist (CS012-13). This report presents independent research funded by the NIHR. The views expressed are those of the authors and not necessarily those of the UK National Health Service, the NIHR or the UK Department of Health.

## Supporting information

**S1 Table.** Table List of parameters and values used in simulations.

| Parameter | Description | Values used |
| --- | --- | --- |
| $V_{FRC}$ | Total lung and mouth cavity volume at rest. | 3 L |
| $V_D$ | Total volume of conducting airways. | 0.12 L |
| $V_{mouth}$ | Total volume of mouth cavity. | 0.05 L |
| $V_T$ | Tidal volume. | 1L |
| $\tau$ | Breath time (inhalation or exhalation). | 2.5 s |
| $V_{duct}/V_{acin}$ | Proportion of acinar gas volume in ducts. | 0.2 [47] |
| $\lambda_{cond}$ | Airway length scaling in mean-path conducting branches. | 0.794 [5, 48] |
| $\lambda_{acin}$ | Duct length scaling in acinar branches. | 0.93 [47, 49] |
| $LD_{cond}$ | Ratio of length to diameter in mean-path conducting branches. | 3 [6, 50] |
| $LD_{acin}$ | Ratio of length to diameter in acinar branches. | 2.3 [47] |
| $N^{acin} + 1$ | Number of acinar airway generations. | 9 [47] |
| $\Phi_j$ | Density of alveolar sacs.<br>in acinar generations.<br>$j = N_z^{cond} \dots N^{tot}$. | 0 if $j < N_z^{cond}$<br>0.2 if $j = N_z^{cond}$<br>0.4 if $j = N_z^{cond} + 1$ [47]<br>0.7 if $j = N_z^{cond} + 2$<br>1 if $j > N_z^{cond} + 2$ |
| $\phi$ | Phenomenological parameter, see eq. (4). | 0.5 [12] |
| $K_{lung}$ | Elasticity of the whole lung. | 5 cm H<sub>2</sub>O L<sup>-1</sup> [51, 52] |
| $R_{acin}$ | Cumulative resistance of acini. | 0.2 cm H<sub>2</sub>O s L<sup>-1</sup> [51, 53] |
| $R_{UA}$ | Resistance of the upper airway. | 0.6 cm H<sub>2</sub>O s L<sup>-1</sup> [51, 53] |
| $\mu$ | Air viscosity at 37°C. | $1.93 \times 10^{-7}$ cm H<sub>2</sub>O s |
| $\{d, l\}$<br>Diameters<br>and lengths<br>of proximal<br>bronchi<br>(cm) | Trachea<br>Right Main Bronchus<br>Left Main Bronchus<br>Right Intermediate Bronchus<br>Right Lower Lobar Bronchus<br>Left Lower Lobar Bronchus | {1.6, 10}<br>{1.11, 2.2}<br>{1.2, 5} [6]<br>{0.89, 2.6}<br>{0.64, 0.8}<br>{0.8, 1.1} |

**S1 Figure.**
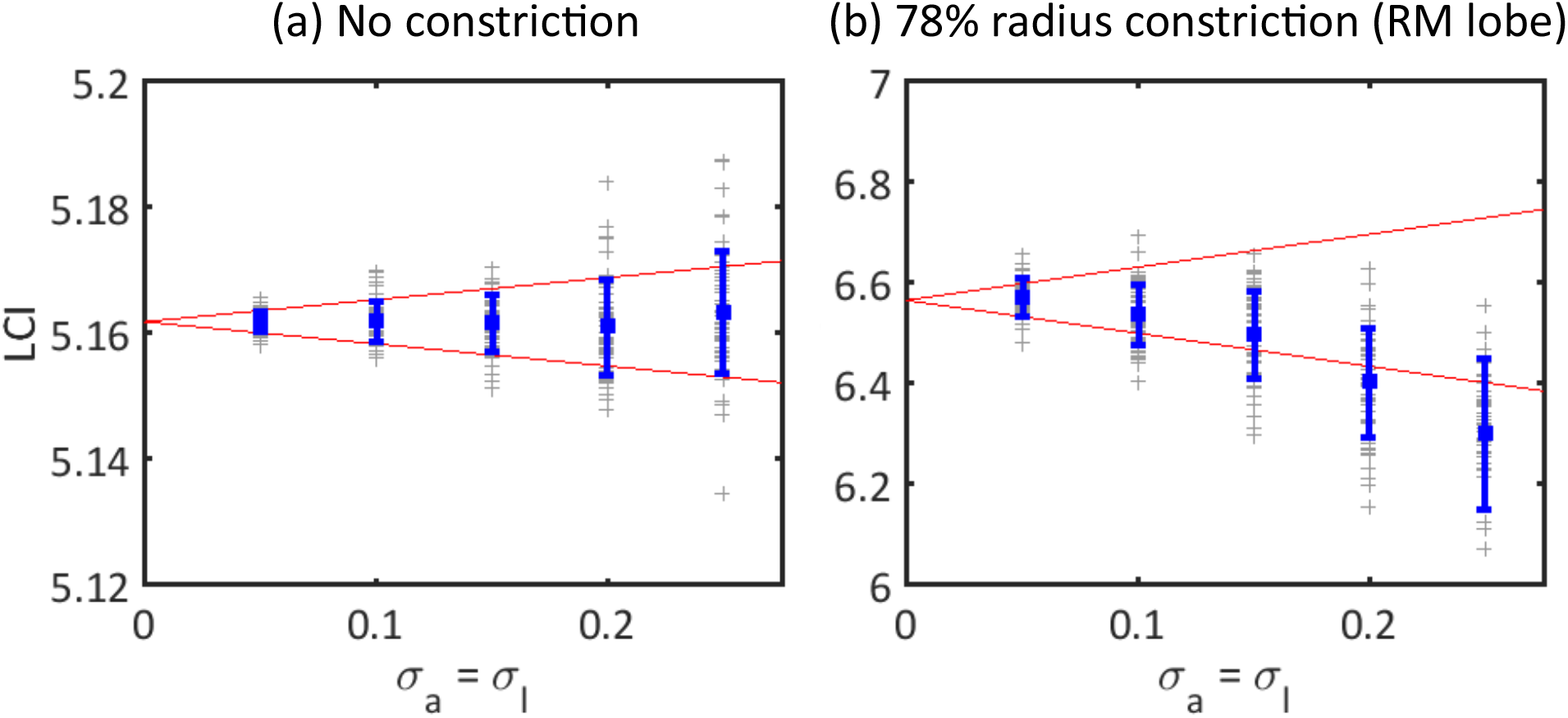
Comparison of model P prediction of LCI variance with Monte Carlo (non-perturbative) simulations. In both cases airway perturbations are assumed independently normally distributed with same coefficient of variation in area and length (*σ_a_* = *σ_l_*) and variance in elasticity is not considered *σ_K_* = 0. Perturbations are only drawn for airways down to and including Strahler order 14, with the remaining generations assumed to be perfectly symmetric (as in model M). For the Monte Carlo prediction, the normal distribution of perturbations is truncated to prevent unphysical behaviour and preserve symmetry such that 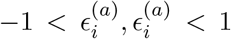. Error bars indicate mean ± one s.d. of model outputs and red lines show the prediction of mean ± one s.d. from model P. Crosses mark the results of individual realisations in the Monte Carlo algorithm. The results are shown for the case when perturbations are applied to (a) the healthy model M, and (b) model M with severe constrictions in the proximal airways of the RM lobe. Good agreement for the predicted variance is observed in both cases up to *σ_a_* = *σ_l_* = 0.25, however in the constricted case heterogeneity tends to result in a lower mean LCI, which is not captured by model P.

**S2 Figure.**
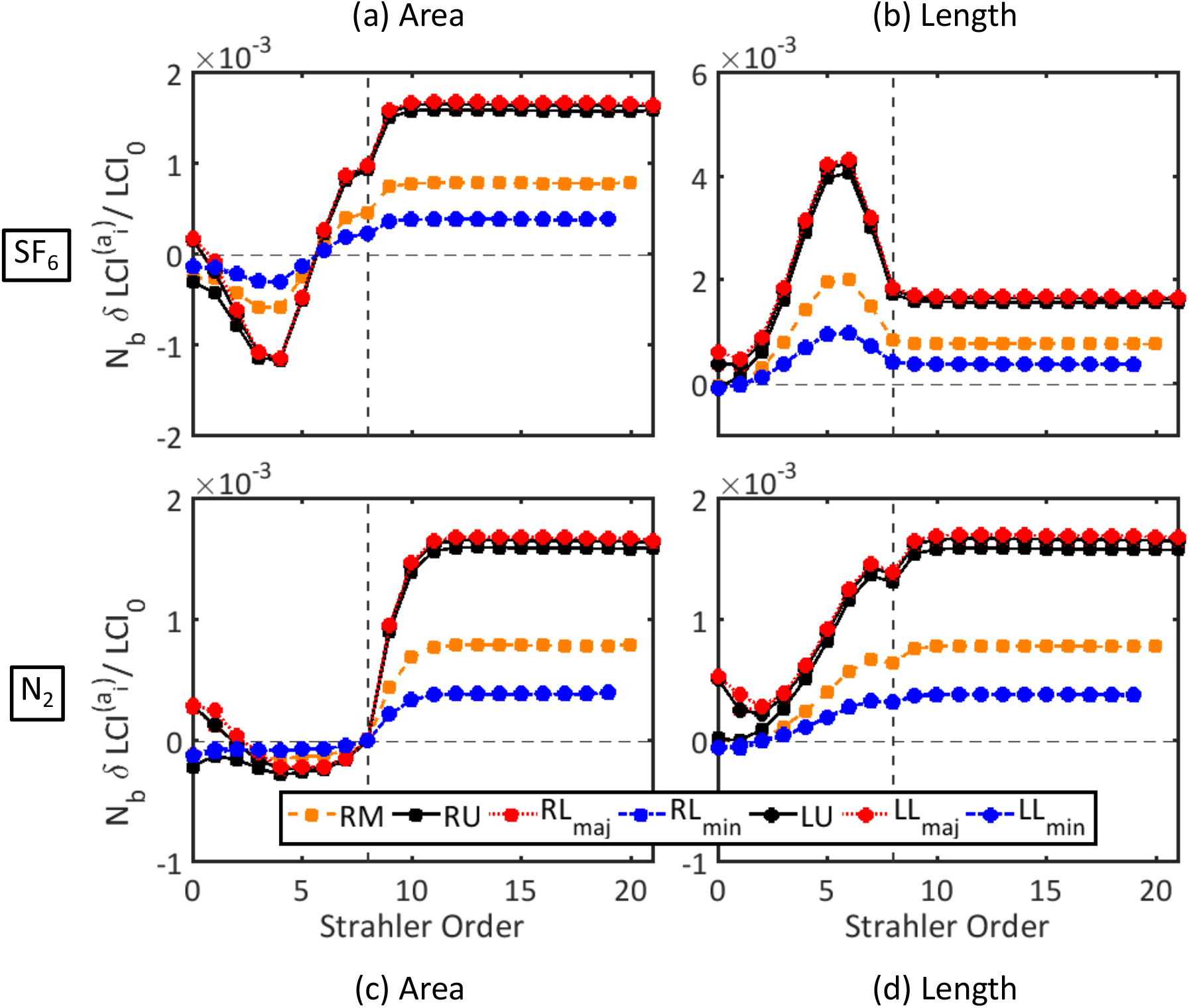
Linear fractional change in LCI (computed for model P) due to a single perturbation in area ((a) and (c)) and length ((b) and (d)) scaled by number of branches in that generation *N_b_*. The airway generation is plotted in terms of its Strahler order (*i.e*. its generation counting up from zero at the bottom of the tree). The vertical dashed line indicate the terminal bronchiole separating the acinar (Strahler orders 0-8) and conducting (>9) generations. (a)-(b) Healthy lung model (no constrictions) using SF_6_ (molecular diffusivity 0.105cm^2^ s^-1^). (c)-(d) Healthy lung model using N_2_ (molecular diffusivity 0.225cm^2^ s^-1^). Coloured symbols distinguish perturbations in the seven lobar regions. The LCI sensitivities in the conducting region (right of the vertical dashed line) are approximately identical for area and length perturbations in both cases, as this is a response to the increase in dead-space volume. For SF_6_ the sensitivities in the acinar region (left of the vertical dashed line) are inverted for length and area perturbations, most notably around the diffusion front (approximately Strahler order 4). Thus LCI is sensitive to geometry changes that affect diffusion in the acinus when using the less diffusive SF_6_, but not N_2_.

**S3 Figure.**
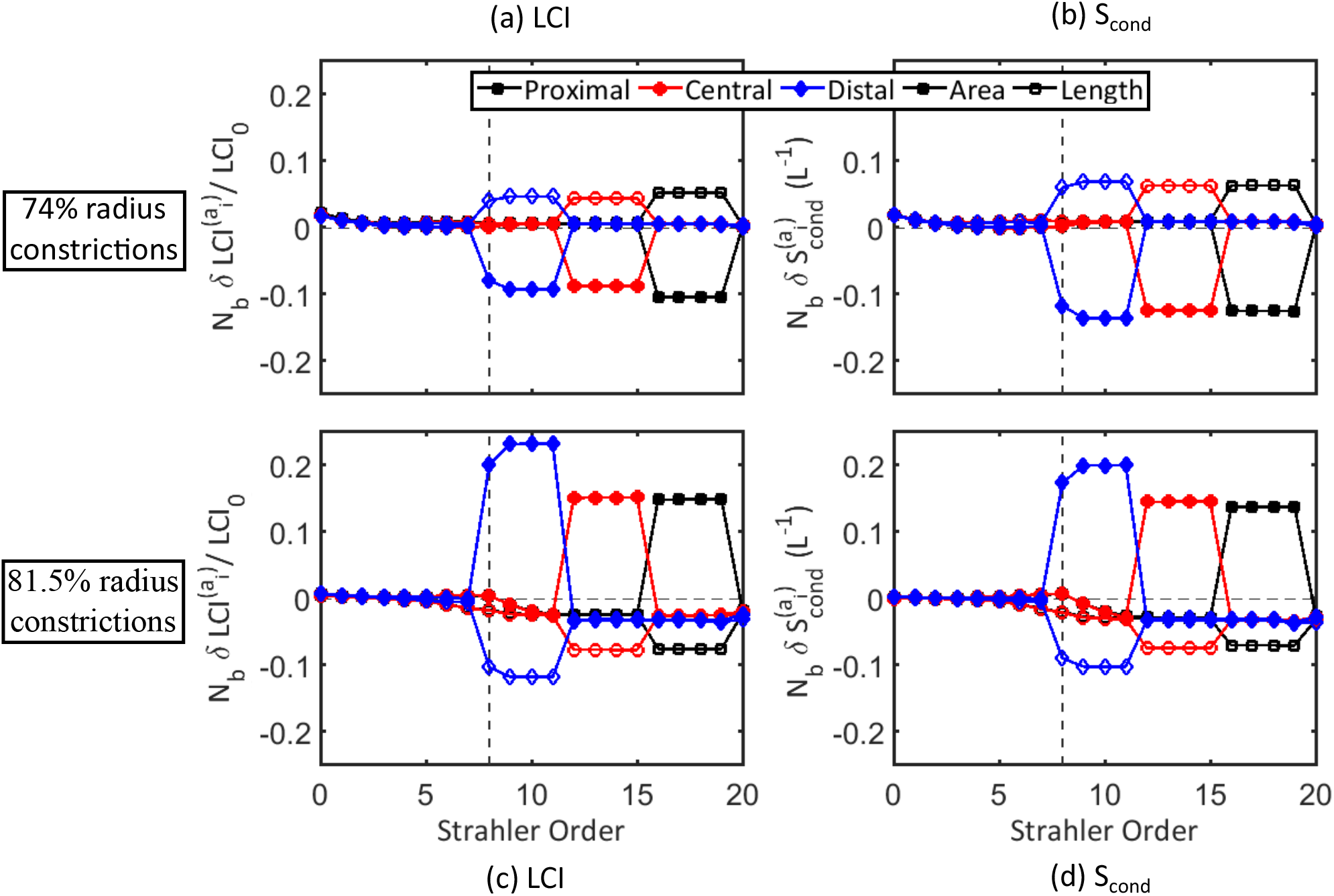
Linear sensitivities (computed for model P) to geometry perturbations in a simulations with (a)-(b) 74% and (c)-(d) 81.5% constrictions in radius to the RM lobe. Scaled sensitivities (as in S2 Figure) w.r.t. area (filled markers) and length (open markers) of the airways are shown for the RM lobe only for fractional LCI change and absolute change in *S*_cond_. Results were plotted for different depths of constriction: proximal (Strahler orders 16-19, black squares), central (Strahler orders 12-15, red circles) and distal (Strahler orders 8-11, blue diamonds). The sensitivities are scaled by the number of airways in the corresponding Strahler order of the RM lobe. The scaled sensitivities are much larger in the constricted airways, as the response is most sensitive to their resistance (note the difference in scale to S2 Figure). Since airway resistance scales as length/area^2^, the area sensitivities are approximately a factor –2 of the length sensitivities. The sign of the sensitivities changes between the two constriction strengths because they lie either side of the maximum values of LCI and *S*_cond_ in Figure 4(a) and (b).

**S1 Video** Inert gas concentration on model M lung network for various constriction strengths to the central airways of the RM lobe (% reduction in radius as shown). Vertical direction is the distance from the mouth, while horizontal distances have no physical meaning and are set for visibility. Time scale 1:4 (each second of video corresponds to 4 seconds of washout).

**S2 Video** Inert gas concentration on model M lung network with randomly distributed constrictions of various magnitudes in the central airways (% reduction in radius as shown). Vertical direction is the distance from the mouth, while horizontal distances have no physical meaning and are set for visibility.. Time scale 1:4 (each second of video corresponds to 4 seconds of washout).

**S1 File** Supplementary text containing further details of the methodology.

